# Selective transcriptomic recovery by (*2R,6R*)-hydroxynorketamine in opioid-abstinent mice: Machine learning identifies predictive biomarkers

**DOI:** 10.1101/2025.06.04.657935

**Authors:** Anna Onisiforou, Morfeas Koumas, Andria Michael, Panos Zanos

## Abstract

**Background:** Opioid abstinence induces persistent emotional disturbances and widespread neuroplastic changes, in areas of the brain including the hippocampus. Although ketamine and its metabolite (*2R,6R*)-hydroxynorketamine (HNK) show potential in reversing opioid abstinence-related deficits in rodents, the molecular mechanisms underlying their efficacy remain poorly understood.

**Methods:** Male C57BL/6J mice underwent a 3-week opioid abstinence paradigm, followed by a single (2R,6R)-HNK (10 mg/kg, i.p.) or saline injection on day 28. Sucrose and social preference tests were used to assess behavioral deficits. We conducted RNA sequencing of ventral hippocampal tissue from these mice, followed by differential gene expression and functional enrichment analyses. Additionally, Random Forest machine was applied to identify predictive differentially expressed genes (DEGs) associated with (*2R,6R*)-HNK treatment response.

**Results:** Transcriptomic analysis identified 206 DEGs in morphine-abstinent mice without treatment compared to opioid-naïve controls (MOR-SAL *vs.* SAL-SAL), implicating altered immune signaling, synaptic function, and structural plasticity. Comparison of opioid-abstinent mice treated with (*2R,6R*)-HNK to opioid-naive controls (MOR-HNK *vs.* SAL-SAL) revealed 186 residual DEGs, enriched for Th17-mediated immune and fear regulation pathways, suggesting a persistent intermediate molecular phenotype despite normalized behavioural scores. DEGs overlap analysis between MOR-HNK *vs*. MOR-SAL and MOR-SAL *vs*. SAL-SAL indicated that (*2R,6R*)-HNK treatment reversed 55 DEGs in opioid-abstinent mice, including Transthyretin (*Ttr*) and T-cell surface glycoprotein (*Cd5*) expression levels. Machine learning identified interleukin 1 receptor accessory protein-like 1 (*Il1rapl1)* and cytotoxic T lymphocyte-associated protein 2 beta (*Ctla2b)* as top predictors of (*2R,6R*)-HNK’s treatment response. Notably, while (2R,6R)-HNK induces transcriptional changes in opioid-naive mice (SAL-HNK), it does not affect behavior compared to untreated controls (SAL-SAL). In contrast, its therapeutic effects are evident in morphine-abstinent mice (MOR-HNK), highlighting its context-dependent efficacy.

**Conclusion:** (2R,6R)-HNK promotes both transcriptional and behavioral recovery in opioid-abstinent mice, reversing key gene expression changes. However, persistent dysregulation of neuroimmune and emotion-related pathways suggests an intermediate molecular state, reflecting ongoing recovery.

## 1. Introduction

Opioid addiction, also referred to as opioid use disorder (OUD), is a chronic neuropsychiatric condition that affects approximately 27 million people worldwide. It is characterized by a high risk of relapse and contributes significantly to global mortality, with approximately 480,000 deaths annually, primarily due to overdoses ^1^. Acknowledging the significance of the crisis, the U.S. officially designated it a national emergency and public health epidemic in 2017 ^2^. Additionally, opioid-related fatalities have been rising in Europe and other parts of the world ^3^.

Prolonged abstinence following chronic opioid use is associated with mood disturbances, such as anhedonia (a diminished ability to experience pleasure) ^4^, heightened anxiety ^5–7^, and social withdrawal ^8^. The emotional distress experienced during extended abstinence, coupled with significant cognitive impairments ^9^, shares similarities with mood disorders ^7,10^. This comorbidity poses a challenge for effectively treating OUD, as it is linked to prolonged illness, a worse prognosis, and a greater likelihood of relapse ^4,11^, with relapse rates reaching up to 91% ^12^.

Stress exposure during opioid abstinence is recognized as a key factor that contributes to relapse ^13^. Both acute and repeated stress have been identified as significant risk factors for opioid misuse and relapse ^14^. Traditional antidepressants, which primarily target monoamine pathways, have shown some effectiveness in improving patient retention in medication-assisted treatment programs ^15^. However, they offer limited success in alleviating negative affect or completely preventing relapse among opioid-abstinent individuals ^16,17^. Similarly, opioid substitution therapies have been crucial in reducing cravings, mitigating abstinence symptoms, and extending periods of abstinence ^18^. Despite their benefits, these medications, often requiring long-term administration, are not entirely effective in addressing negative affect or preventing relapse ^18^. Research suggests that between 20% and 57% of individuals undergoing methadone maintenance therapy resume opioid use within six months, with relapse rates increasing over time ^19,20^. Additionally, depression remains a common co-occurring condition ^21^. While buprenorphine has been shown to reduce depressive symptoms in OUD patients, many still struggle with significant mood disturbances ^22,23^. The fact that a majority relapse within six months highlights the critical need for treatments that address the emotional challenges associated with OUD ^24^. Furthermore, up to 56% of individuals require additional antidepressants or other psychotropic medications during prolonged abstinence ^25–30^. Consequently, the development of novel, fast-acting treatments is essential for improving OUD outcomes and reducing relapse rates.

Emerging evidence suggests that (*R,S*)-ketamine, known for its rapid antidepressant effects, may aid in maintaining opioid abstinence and reducing cravings, thus possibly preventing relapse ^31–35^. Preclinical studies align with clinical findings, showing that both racemic ketamine ^36^ and the (*R*)-ketamine enantiomer ^37^ mitigate somatic withdrawal symptoms in opioid-dependent animal models. Despite these promising results, the feasibility of using ketamine as a daily treatment for OUD is limited due to its dissociative side effects and potential for abuse ^38^. Recent preclinical studies suggest that ketamine’s metabolite (*2R,6R*)-HNK reduces acute withdrawal-induced somatic symptoms and prevents reinstatement of opioid-seeking behaviors following extinction. Additionally, this metabolite has been shown to decrease anxiety-like behaviors in acute opioid-abstinent mice ^39^. In a recent study, we showed that (*2R,6R*)-HNK reverses opioid-induced conditioning and withdrawal symptoms in mice ^40^. We also showed that (*2R,6R*)-HNK could alleviate affective behaviors during abstinence via a GluN2A-NMDA receptor-dependent restoration of cortical activity ^40^.

Despite the behavioral promise of (2R,6R)-HNK in OUD models, its underlying molecular mechanisms remain largely uncharacterized, particularly in brain regions like the hippocampus, which are critical for affective regulation and memory and are profoundly affected during protracted opioid abstinence. To address this, we analyzed hippocampal gene expression across four treatment conditions: saline in opioid-naive controls (SAL-SAL), (*2R,6R*)-HNK in opioid-naive controls (SAL-HNK), saline in morphine-abstinent mice (MOR-SAL), and (*2R,6R*)-HNK in morphine-abstinent mice (MOR-HNK) (see **Figure 1**). Comparisons of MOR-SAL *vs.* SAL-SAL revealed protracted opioid abstinence-induced transcriptomic alterations, while MOR-HNK *vs.* MOR-SAL identified gene expression changes associated with (*2R,6R*)-HNK-mediated reversal. Comparisons of MOR-HNK *vs.* SAL-SAL assessed the extent to which (*2R,6R*)-HNK normalized morphine-altered gene expression toward a baseline profile. Additionally, SAL-HNK *vs.* SAL-SAL comparisons allowed us to evaluate the transcriptional effects of (*2R,6R*)-HNK in the absence of prior drug exposure, distinguishing general drug-induced changes from those that are context-dependent. This approach enabled a systematic mapping of the molecular landscape of opioid-induced dysfunction and the potential therapeutic modulation by (*2R,6R*)-HNK.

**Figure 1:**
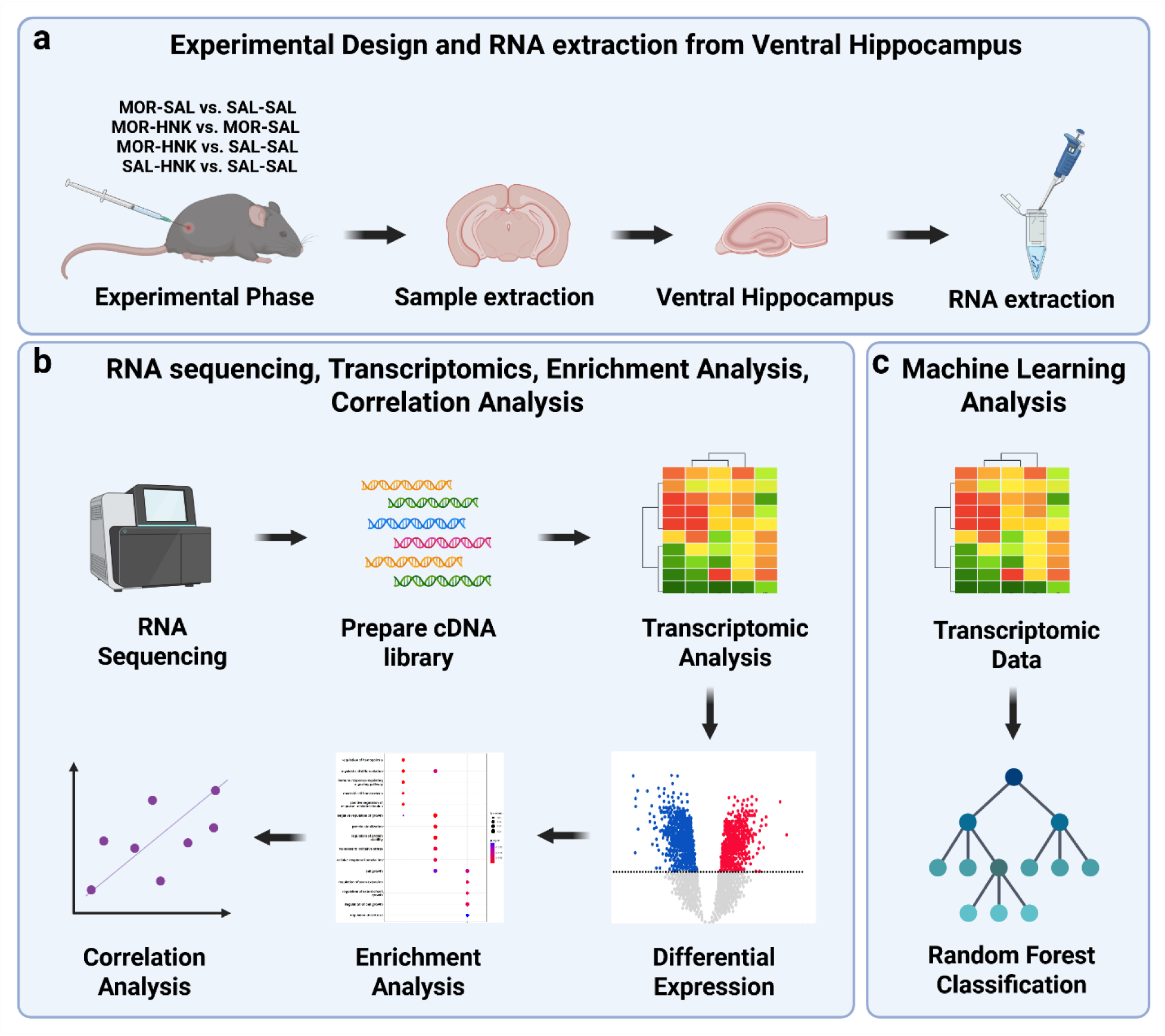
Overview of Experimental Workflow for Transcriptomic and Machine Learning Analysis of (2R,6R)-HNK Effects in Morphine-Abstinent Mice. **(a)** Schematic of the experimental design, including treatment groups and ventral hippocampus RNA extraction. **(b)** RNA sequencing workflow including transcriptomic profiling, enrichment analysis, and behavioral correlation. **(c)** Machine learning analysis using Random Forest classification to identify predictive gene signatures.

## 2. Methods

### 2.1 Animals

Male C57BL/6J mice (8 weeks old, Jackson laboratories) were kept in a temperature and humidity-controlled environment and maintained in a 12:12 hr light/dark cycle (lights on at 7:00AM). Mice were singly housed for the study described due to increased aggression induced by opioid administration and were randomly assigned to treatment groups. All experimental procedures were approved by the Cyprus National Committee for Animal Welfare and were conducted in full accordance with the National Guides for the Care and Use of Laboratory Animals and reported according to ARRIVE guidelines.

### 2.2 Drugs

(*2R,6R*)-HNK hydrochloride was synthesized and characterized at the National Center for Advancing Translational Sciences (NCATS, NIH, U.S.A.). Absolute and relative stereochemistry for (*2R,6R*)-HNK was confirmed by small molecule x-ray crystallography ^41^. Morphine hydrochloride was purchased from Cayman Europe and was injected subcutaneously (s.c.) at different doses of 20, 40, 60, 80, 100 mg/kg. For the intraperitoneal injections, (*2R,6R*)-hydroxynorketamine (HNK) was dissolved in 0.9% saline and given in a volume of 10mg/kg.

### 2.3 Chronic escalating-dose morphine administration and protracted abstinence paradigm

Protracted opioid abstinent individuals experience various emotional disturbances such as depression, anxiety along with other cognitive impairments, which can serve as a trigger for relapse. In this study we aimed at assessing ketamine’s metabolite (*2R,6R*)-HNK, in reversing the assessed maladaptive behaviours observed in protracted opioid abstinent mice. To assess this, we used the chronic escalating-dose morphine administration paradigm, previously published by our group ^42,43^ and others, including Prof. Brigitte Kieffer’s group ^44–46^. This protocol has demonstrated predictive validity through behavioural tests conducted during protracted abstinence. These studies reliably replicate the emotional deficits observed in humans abstaining from chronic opioid use. Moreover, using this protocol in mice, functional connectivity analyses with fMRI have revealed durable, brain-wide alterations and identified specific biomarkers (e.g., retrosplenial cortex and amygdala FC) also observed in humans, underscoring the translatability and relevance of this model for OUD research ^46^.

Briefly, male C57BL/6J mice (n=8 per group) were administered with escalating doses of morphine (20, 40, 60, 80, 100 mg/kg) or saline (s.c.) twice daily, for a total of 5 days. On day 6, a single 100mg/kg injection of either morphine or saline was used. To induce protracted abstinence, mice were allowed to withdraw in a drug-free environment for a total of 28 days. On Day 28 of abstinence, a single injection of (*2R,6R*)-HNK (10mg/kg; i.p.) or saline was administered. Mice were tested for sucrose preference overnight and for social preference on Day 29, as we previously published ^41,47^. On Day 29 of abstinence, mice were euthanized via decapitation and brains were excised, and the ventral hippocampi was immediately dissected and stored in −80°C for preservation.

### 2.4 Behavioral data statistical analysis

Experimentation and analysis were performed in a manner blind to treatment assignments. Mice were randomly assigned to treatment groups. Statistical analyses were performed using GraphPad Prism software *v*10.1.2. Significance was assigned at *p* < 0.05. Normality and equal variances between group samples were assessed using the Kolmogorov-Smirnov and Brown-Forsythe tests, respectively. Sucrose and social preference data were analyzed using two-way analysis of variance (two-way ANOVA). ANOVAs were followed by a Holm-Šídák post-hoc comparison when both normality and equal variances were confirmed. Results are presented as mean ± SEM.

### 2.5 RNA extraction

RNA extraction was performed using the RNeasy Mini Kit (Qiagen, Cat.No. 74106), according to the manufacturer’s instructions. Briefly, ventral hippocampi (∼15-30mg) were homogenized in 350μl lysis solution (Buffer RLT) and centrifuged at maximum speed for 3mins. Equal volume (350µl) of 70% ethanol was added to the lysate and the total of 700µl was transferred to the mini spin column and centrifuged for 15secs at ≥8000 x g. The supernatant was discarded and 700µl of Buffer RW1 was added to the column and centrifuged for 15secs at ≥8000 x g. The supernatant was again discarded and 500µl of Buffer RPE was added to the column and centrifuged for 15secs at ≥8000 x g. A further 500µl of Buffer RPE was added to the column and centrifuged for 2mins at ≥ 8000 x g. The spin column was then placed in a new 1.5ml collection tube and 30µl of RNase-free water was added. To elute the RNA, the spin column and collection tube were them centrifuged for 2mins at ≥8000 x g. RNA concentration was measured with a Nanodrop (NanoPhotometer N60/N50, Implen).

### 2.6 RNA sequencing and analysis

RNA sequencing was performed by Novogene Europe. Messenger RNA was purified from total RNA using poly-T oligo-attached magnetic beads. After fragmentation, the first strand cDNA was synthesized using random hexamer primers, followed by the second strand cDNA synthesis using either dUTP for directional library or dTTP for non-directional library ^48^. For the non-directional library, it was ready after end repair, A-tailing, adapter ligation, size selection, amplification, and purification. For the directional library, it was ready after end repair, A-tailing, adapter ligation, size selection, USER enzyme digestion, amplification, and purification. The library was checked with Qubit and real-time PCR for quantification and bioanalyzer for size distribution detection. Quantified libraries were pooled and sequenced on the Illumina NovaSeq 6000, according to effective library concentration and data amount. Raw data (raw reads) of fastq format were firstly processed through in-house perl scripts by Novogene Europe. In this step, clean data (clean reads) were obtained by removing reads containing adapter, reads containing ploy-N and low-quality reads from raw data. At the same time, Q20, Q30 and GC content the clean data were calculated. All the downstream analyses were based on the clean data with high quality.

The clean raw sequence data (FASTQ files) were mapped to the mouse reference genome (mm10) using HISAT2 ^49^ (Galaxy Version 2.2.1+galaxy1), generating BAM files. Unique gene hit counts were quantified using FeatureCounts (Galaxy Version 2.0.8+galaxy0), with the minimum mapping quality per read set to 10, meaning only reads with a mapping quality score of at least 10 were counted. Differential gene expression analysis was performed using DESeq2 to compare MOR-SAL *vs*. SAL-SAL, SAL-HNK *vs*. SAL-SAL, MOR-HNK *vs*. SAL-SAL and MOR-HNK *vs*. MOR-SAL. The Wald test was used to compute *p*-values and log2 fold changes. Genes were considered significantly differentially expressed with a *p*-value < 0.05 and an absolute log2 fold change (|log2FC| > 0.5). A pre-filtering step was applied to remove genes with low read counts (≤10 across all samples) before differential expression analysis.

### 2.7 Comparison of oppositely regulated genes between MOR-SAL vs. SAL-SAL and MOR-HNK vs. MOR-SAL

To identify genes that were oppositely regulated between the two comparisons, DESeq2 results were merged by gene symbol. Genes were selected if they showed opposite regulation in the two conditions, meaning they were upregulated in one group and downregulated in the other.

The magnitude of expression difference was calculated as:

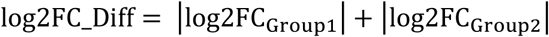

ensuring that only genes with complete expression switching were considered.

The sign of log2FC_Diff was assigned based on the dominant regulation, where negative values indicate overall downregulation, and positive values indicate overall upregulation.

### 2.8 Functional enrichment analysis

To validate the biological significance of DEGs in MOR-SAL *vs*. SAL-SAL, SAL-HNK *vs.* SAL-SAL, MOR-HNK *vs*. SAL-SAL and MOR-HNK *vs*. MOR-SAL comparisons, Gene Ontology (GO) Biological processes and Kyoto Encyclopedia of Genes and Genomes (KEGG) pathway enrichment analyses were performed using the ClueGO app ^50^ in Cytoscape. Only statistically significant pathways and GO terms with an adjusted *p*-value ≤ 0.05, corrected using the Bonferroni step-down method, were retained.

### 2.9 Gene expression and behavioral correlation analysis

To investigate whether transcriptional reversal by (*2R,6R*)-HNK predicts behavioral recovery following chronic morphine exposure, we focused our analysis on the top DEGs that showed opposite regulation in the MOR-SAL *vs*. SAL-SAL and MOR-HNK *vs*. MOR-SAL comparisons (see Section 2.7). Using the mapped read counts from each sample in the MOR-HNK group (8 samples), we extracted and isolated the top reversed DEGs for each sample. Gene expression data were log2-transformed prior to correlation analysis to normalize variance across samples. Behavioral outcomes were measured using sucrose preference and social preference tests. Correlations between normalized gene expression levels and individual behavioral scores were assessed using Spearman’s rank correlation coefficient (ρ) within the MOR-HNK group. Unadjusted *p*-values were calculated, and false discovery rate (FDR) correction was applied using the Benjamini-Hochberg procedure.

### 2.10 Application of random forest machine learning to identify predictive DEGs associated with treatment response to (*2R,6R*)-HNK

To identify genes most predictive of treatment response, we applied a Random Forest classification model using the 200 DEGs identified between MOR-HNK *vs.* MOR-SAL mice. We selected the Random Forest classifier for its ability to handle high-dimensional transcriptomic data, provide interpretable gene importance rankings, and offer robust internal validation through out-of-bag error (OOB) estimation, making it particularly well-suited for biomarker discovery in small-sample, high-feature RNA-seq datasets. This supervised machine learning approach was used to rank genes based on their contribution to distinguishing individual samples by treatment group, independent of directional reversal. The resulting importance scores identified candidate molecular markers that best separate (*2R,6R*)-HNK-treated animals from morphine-exposed controls (MOR-HNK *vs.* MOR-SAL). We used Random Forest classification with 500 trees and default OOB estimation, which provides an internal cross-validation-based error rate without the need for an external train-test split.

## 3. Results

### 3.1 Comparison of reward and social behaviors across treatment groups

Behavioral assays revealed that MOR-HNK mice exhibited significantly improved sucrose preference (adjusted *p* < 0.0001) and enhanced social preference (adjusted *p* < 0.0001) compared to MOR-SAL mice (**Figure 2a, b**), indicating that (*2R,6R*)-HNK treatment effectively ameliorated morphine-induced deficits in reward and social behavior. No significant differences were observed in either sucrose (adjusted *p* = 0.7172) or social preference (adjusted *p* = 0.8041) (**Figure 2a, b**) between the MOR-HNK *vs.* SAL-SAL mice, indicating that overt behavioral performance in these groups was comparable under these standard assay conditions. Similarly, no significant difference was observed in either sucrose (adjusted *p* = 0.6816) or social preference (adjusted *p* = 0.8041) (**Figure 2a, b**) between the SAL-HNK *vs.* SAL-SAL mice, suggesting that (*2R,6R*)-HNK does not induce overt behavioral alterations in opioid-naive animals under standard assay conditions. In contrast, MOR-SAL mice exhibited significantly reduced sucrose preference (adjusted p < 0.0001) and social preference (adjusted p < 0.0001) compared to SAL-SAL controls (**Figure 2a, b**), confirming that morphine abstinence induced deficits in reward and social behavior.

**Figure 2.**
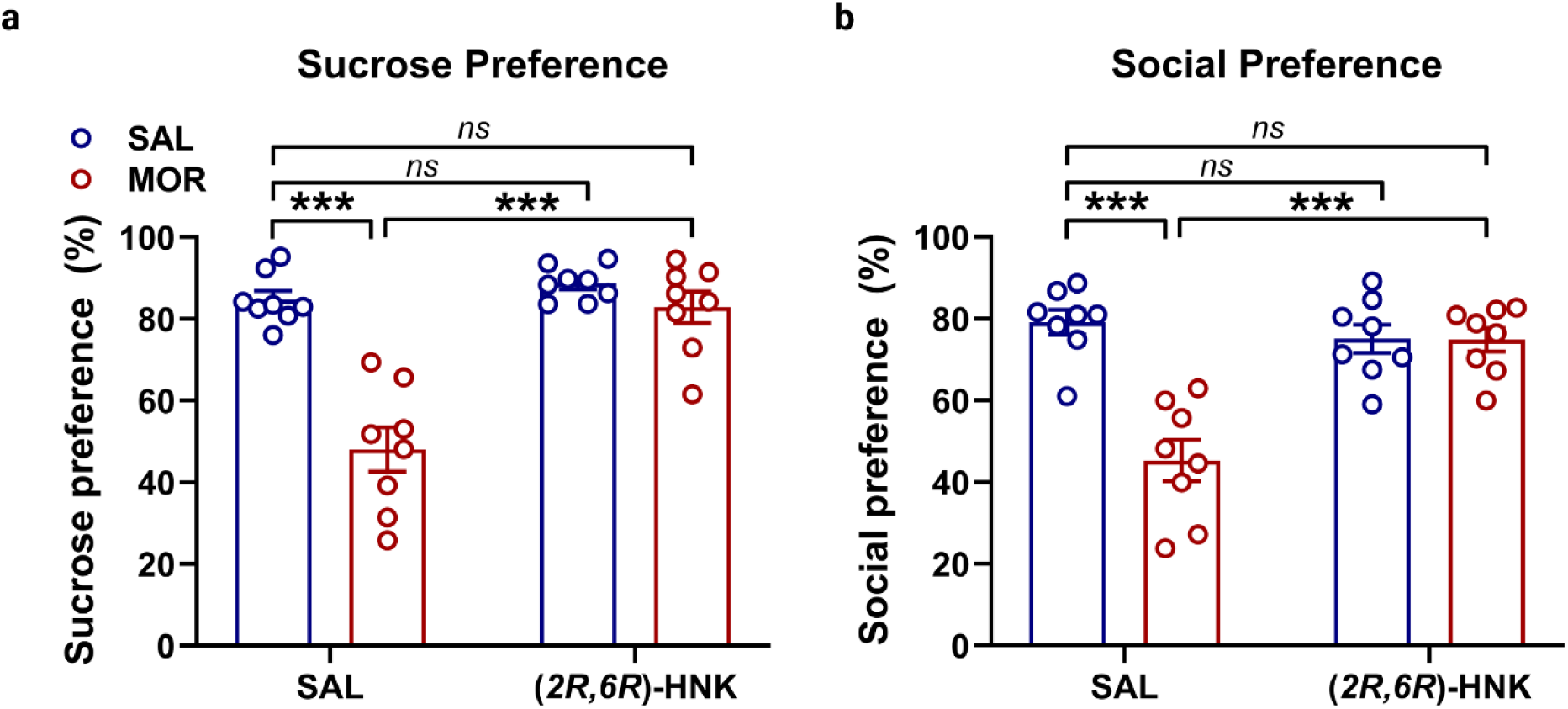
Comparison of Reward and Social Behaviors Across Treatment Groups. **(a)** Sucrose preference scores and **(b)** social preference scores.

### 3.2 Transcriptional alterations induced by morphine in the mouse ventral hippocampus compared to saline-treated control mice

To evaluate the validity of the chronic escalating-dose morphine model, we performed differential expression (DE) analysis comparing MOR-SAL *vs.* SAL-SAL mice, which identified 206 DEGs, including 113 downregulated and 93 upregulated genes, filtered by a *p*-value < 0.05 and absolute log2 fold change (|log2FC| > 0.5) (**Figure 3a**). Enrichment analysis of all 206 DEGs revealed eight statistically significant KEGG pathways, with 12.5% of the terms associated with the morphine addiction functional group (see **Figure 3b**). These results support the model’s relevance for capturing transcriptional changes linked to opioid exposure and addiction-related neuroadaptations. Enrichment analysis of the 93 upregulated DEGs revealed that morphine upregulates the T-helper 17 cell differentiation pathway and the negative regulation of adaptive immune response (**Figure 3c**).

**Figure 3.**
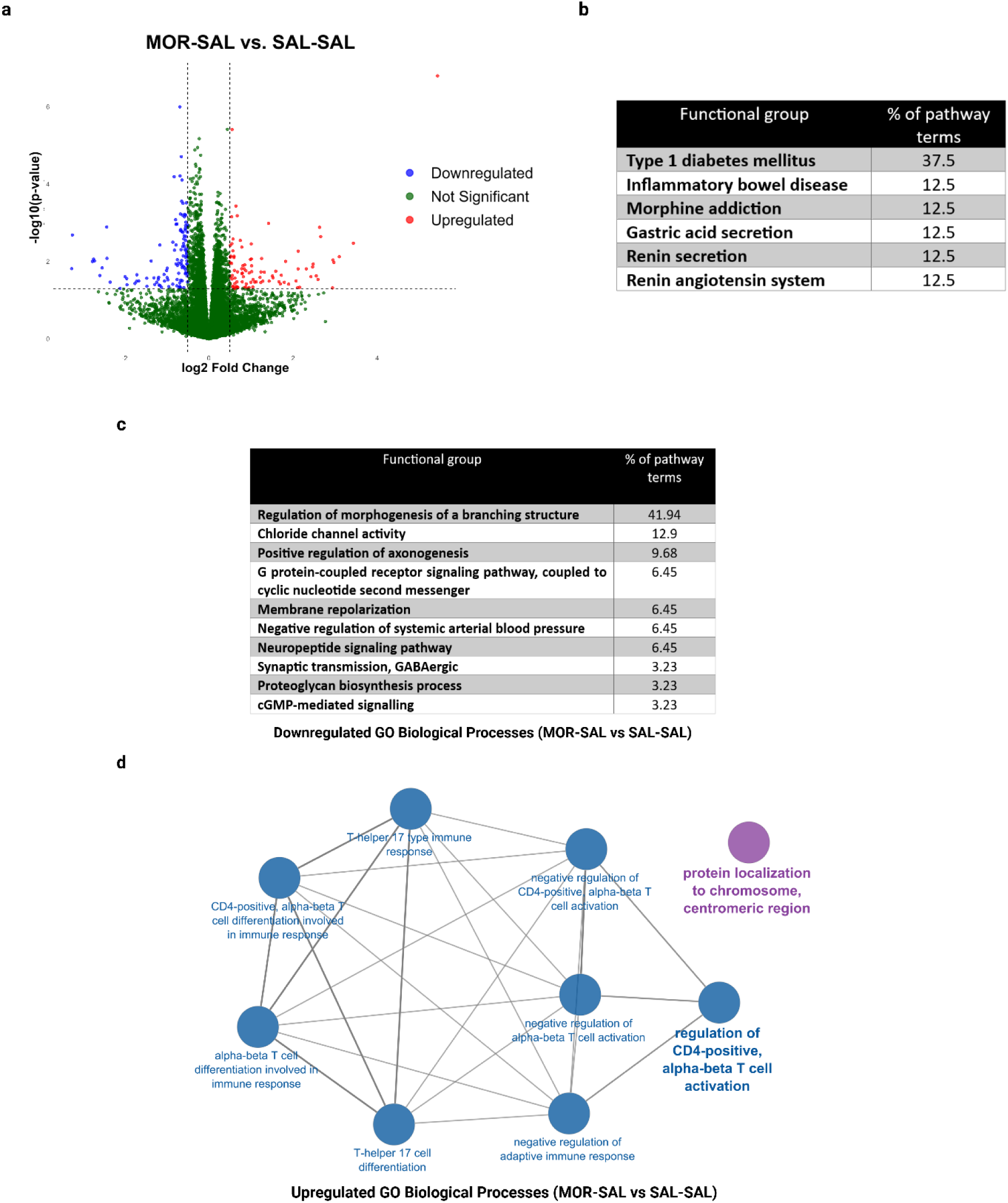
Transcriptomic alterations in the ventral hippocampus induced by chronic morphine exposure. **(a)** Volcano plot showing DE analysis comparing morphine-exposed (MOR-SAL) to saline-treated control mice (SAL-SAL) identified 206 DEGs (p < 0.05, |log₂FC| > 0.5). **(b)** KEGG pathway enrichment of all 206 DEGs revealed eight significantly enriched KEGG pathways. **(c)** GO analysis of the 93 upregulated DEGs. **(d)** GO analysis of 113 downregulated DEGs revealed 31 significantly enriched biological processes across 10 functional groups.

Additionally, enrichment analysis of the 113 downregulated DEGs revealed 31 statistically significant GO biological processes belonging to 10 functional groups (**Figure 3d**). The largest proportion of enriched terms (41.94%) was related to the regulation of morphogenesis of branching structures, suggesting impaired or reduced structural plasticity, including neuronal dendrite and axon formation, following chronic morphine exposure. Other prominent functional groups included chloride channel activity (12.9%), positive regulation of axonogenesis (9.68%), G protein-coupled receptor (GPCR) signaling coupled to cyclic nucleotide second messenger systems (6.45%), and membrane repolarization (6.45%), indicating disruption of ionic homeostasis and synaptic signaling processes. Functional terms associated with neuropeptide signaling (6.45%) and synaptic transmission, specifically GABAergic signaling (3.23%), were also downregulated, suggesting broad suppression of neurotransmission pathways. Additional downregulated processes included the negative regulation of systemic arterial blood pressure (6.45%), proteoglycan biosynthesis (3.23%), and cGMP-mediated signaling (3.23%), which may reflect impairments in vascular and extracellular matrix function critical for neural plasticity and recovery.

### 3.3 Transcriptional alterations induced by (*2R,6R*)-HNK in the mouse ventral hippocampus compared to saline-treated control mice

To evaluate the effects of (*2R,6R*)-HNK in the absence of opioid exposure, DE analysis comparing SAL-HNK *vs.* SAL-SAL mice identified 141 DEGs, filtered using a *p*-value < 0.05 and an absolute log2 fold change (|log2FC| > 0.5) (**Figure 4a**). Of these, 90 DEGs were upregulated and 51 were downregulated. Among the most significant upregulated genes were *Chrna10* (cholinergic receptor nicotinic alpha 10 subunit) and *Rptoros* (regulatory-associated protein of mTOR, complex 1, opposite strand). These genes are involved in cholinergic signaling and mTOR pathway regulation, indicating potential changes in neuronal communication and cellular metabolism. Among the top downregulated genes were *Dbhos* (dopamine beta-hydroxylase, opposite strand) and *Erich4* (glutamate-rich 4). Their decreased expression may reflect alterations in catecholamine biosynthesis and immune function.

**Figure 4.**
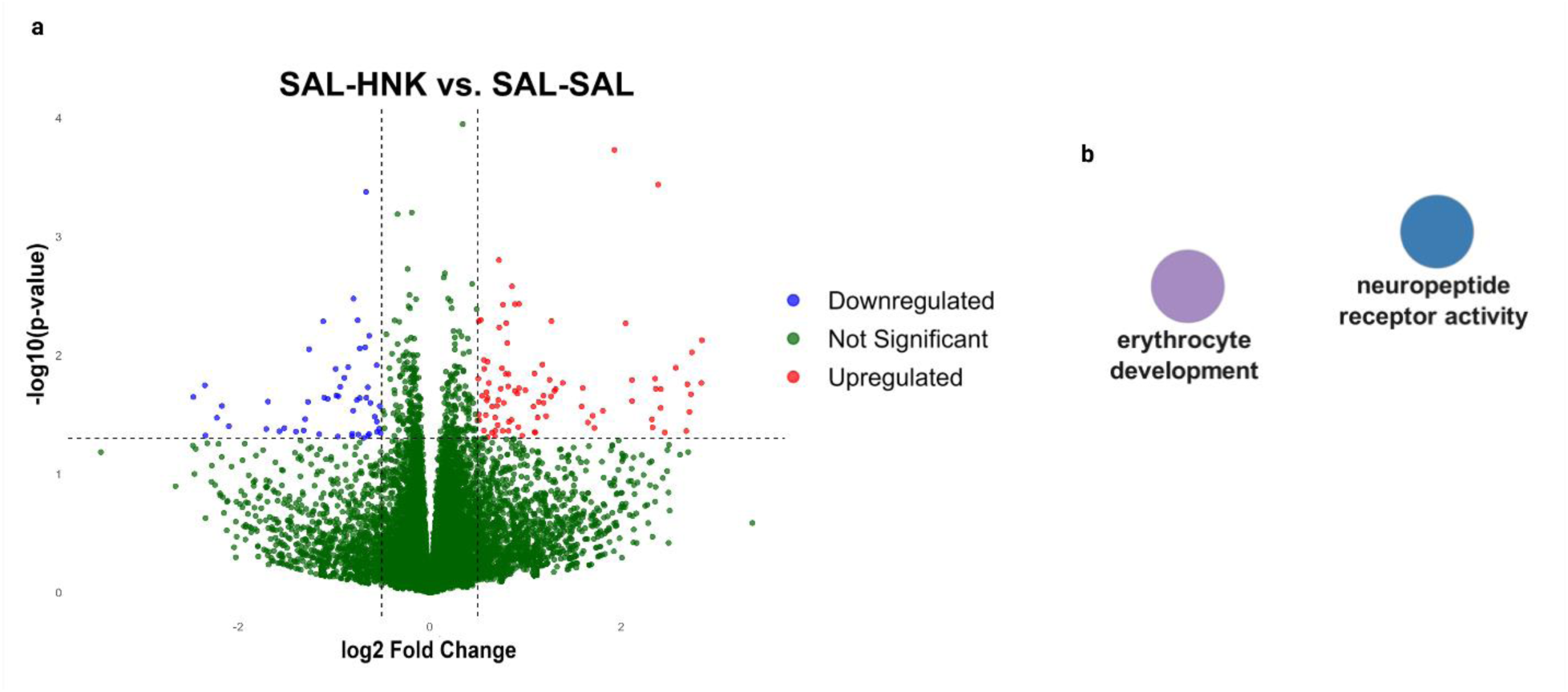
Transcriptomic effects of (2R,6R)-HNK in the absence of morphine exposure. **(a)** Volcano plot showing DEGs between SAL-HNK and SAL-SAL mice, filtered by p-value < 0.05 and |log2 fold change| > 0.5. **(b)** GO enrichment analysis of all DEGs revealed two significantly enriched biological pathways.

GO biological process enrichment analysis performed separately on the upregulated and downregulated DEGs revealed no statistically significant results. However, enrichment analysis of all DEGs identified only two significantly enriched pathways: erythrocyte development and neuropeptide receptor activity (**Figure 4b**), suggesting that while there are measurable transcriptional shifts, their broader functional impact may be limited or context-specific in opioid-naive brains.

### 3.4 (*2R,6R*)-HNK partially reverses morphine-induced transcriptional changes in the ventral hippocampus of morphine-abstinent mice

To determine the ability of (*2R,6R*)-HNK to reverse the morphine-induced transcriptional changes in the chronic escalating-dose morphine mouse model we compared the transcriptomic signature between MOR-SAL *vs*. SAL-SAL and MOR-HNK *vs*. MOR-SAL groups. DE analysis of MOR-HNK *vs.* MOR-SAL (p-value < 0.05, |log2FC| > 0.5) identified 200 DEGs (**Figure 5a**). Venny comparison between MOR-SAL *vs*. SAL-SAL and MOR-HNK *vs*. MOR-SAL revealed 55 common DEGs, 151 DEGs unique to MOR-SAL *vs*. SAL-SAL and 145 DEGs unique to MOR-HNK *vs*. MOR-SAL (**Figure 5b**). To further characterize these 55 common DEGs, we identified genes that exhibited opposite regulation between the two comparisons, that is, genes upregulated in MOR-SAL *vs*. SAL-SAL and downregulated in MOR-HNK *vs*. MOR-SAL, or vice versa. This was achieved by merging DESeq2 results and calculating the sum of absolute log2 fold changes for each gene, ensuring complete expression switching. This analysis confirmed that all 55 common DEGs displayed opposite patterns of regulation, indicative of (*2R,6R*)-HNK’s partial reversal of morphine-induced transcriptional alterations (**Figure 5c**).

**Figure 5:**
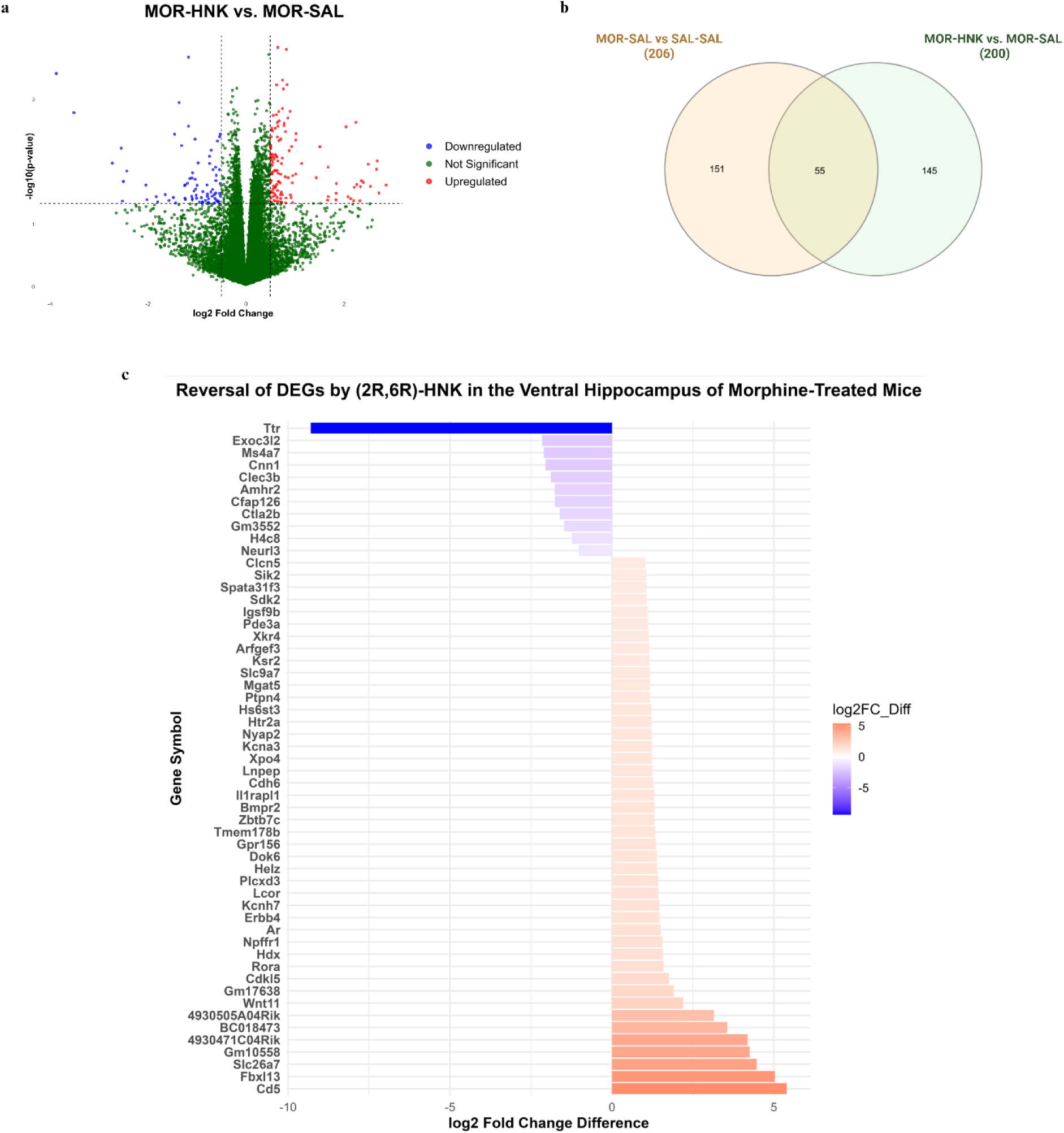
Transcriptomic effects of (2R,6R)-HNK treatment in morphine-exposed mice. **(a)** Volcano plot showing DEGs in the MOR-HNK vs. MOR-SAL comparison (p < 0.05, |log₂FC| > 0.5). **(b)** Venn diagram illustrating the overlap of DEGs between MOR-SAL vs. SAL-SAL and MOR-HNK vs. MOR-SAL comparisons. **(c)** Plot of the 55 overlapping DEGs displaying opposite regulation between MOR-SAL vs. SAL-SAL and MOR-HNK vs. MOR-SAL comparisons.

*Ttr* (transthyretin) expression was significantly upregulated in MOR-SAL mice compared to SAL-SAL controls (log₂FC = +5.4329) and subsequently downregulated in MOR-HNK mice compared to MOR-SAL (log₂FC = –3.8669), resulting in a net reversal log₂FC difference of –9.2998. Similarly, *Cd5* (T-cell surface glycoprotein) expression was significantly downregulated in MOR-SAL mice compared to SAL-SAL (log₂FC = –2.7163) and subsequently upregulated in MOR-HNK mice compared to MOR-SAL (log₂FC = +2.6807), resulting in a net reversal log₂FC difference of +5.3970. *Ttr* is a carrier protein associated with stress and neuroinflammation, and its reversal and downregulation by (2R,6R)-HNK may reflect normalization of stress-induced molecular states. *Cd5*, an immunoregulatory molecule expressed on T cells, was upregulated following HNK treatment, suggesting restoration of immune balance.

### 3.5 Selective association of gene expression reversal with behavioral outcomes

We next examined whether the reversal of morphine abstinence-induced transcriptional changes by (*2R,6R*)-HNK correlated with improvements in behavioral outcomes. Specifically, we assessed whether the robust reversal of *Transthyretin* (*Ttr*, log₂FC = –9.30) and *T-cell surface glycoprotein* (*Cd5*, log₂FC = +5.40) expression levels by (*2R,6R*)-HNK in morphine-exposed mice was associated with performance in sucrose preference and social preference tests, two key behavioral measures of reward and social function.

While *Ttr* was the most strongly reversed DEG following (*2R,6R*)-HNK treatment, exhibiting the greatest downregulation among the identified oppositely regulated genes, its expression levels did not significantly correlate with either social preference (Spearman’s ρ = –0.36, *p* = 0.39) (**Figure 6a**) or sucrose preference (Spearman’s ρ = –0.38, *p* = 0.36) (**Figure 6b**). Similarly, *Cd5*, the most strongly upregulated DEG among the reversal set, also did not significantly correlate with social preference (Spearman’s ρ = –0.21, *p* = 0.62) (**Figure 6c**) or sucrose preference (Spearman’s ρ = –0.16, *p* = 0.71) **(Figure 6d**), suggesting that Cd5 regulation is unlikely to directly contribute to behavioral recovery outcomes. These findings indicate that while (*2R,6R*)-HNK reverses expression of multiple morphine-induced genes, most do not exhibit functional associations with behavioral recovery.

**Figure 6.**
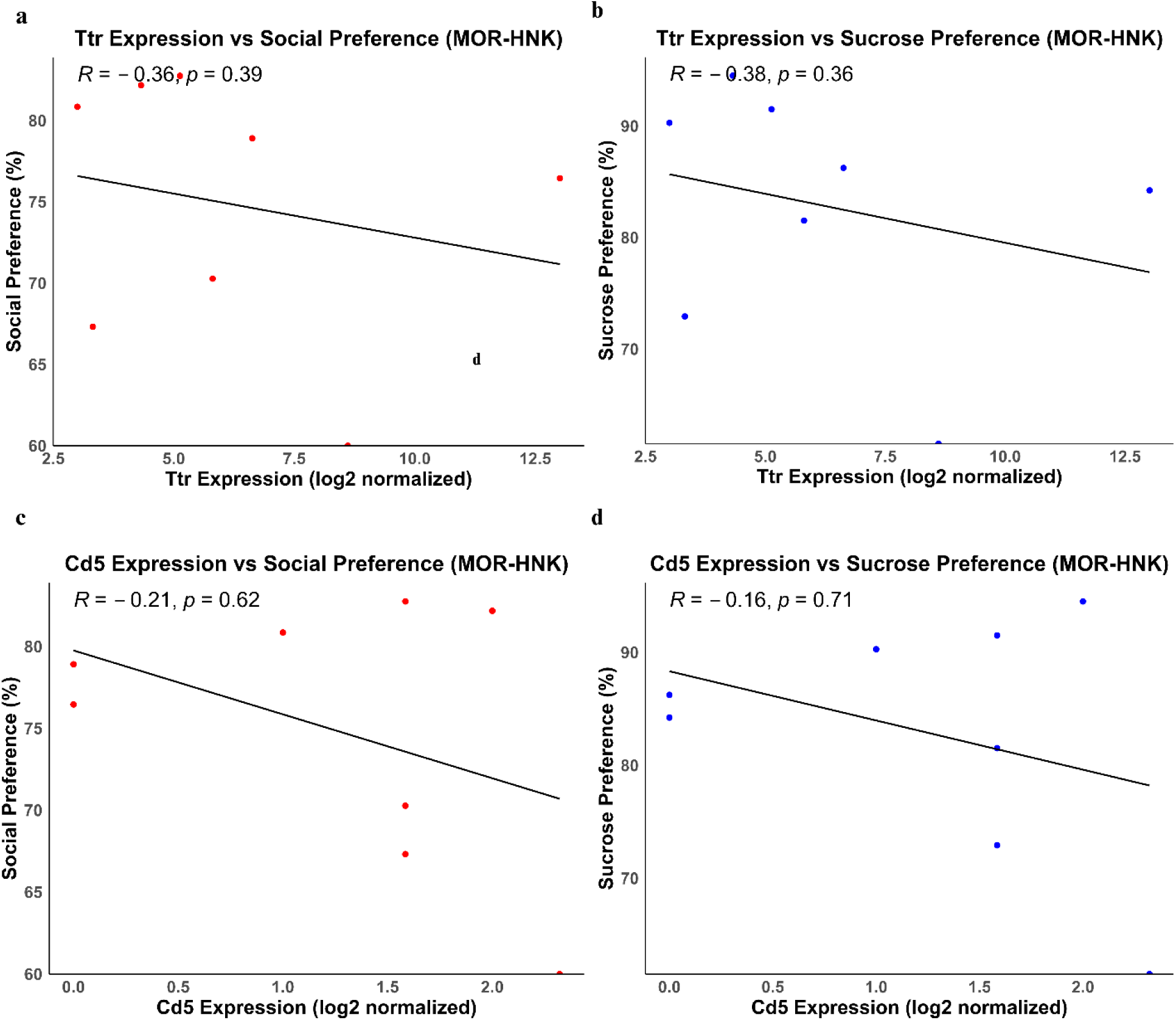
Correlation of reversal gene expression with behavioral performance following (2R,6R)-HNK treatment. **(a)** Correlation between Ttr expression levels and social and **(b)** sucrose preference scores in MOR-HNK mice. **(c)** Correlation between Cd5 expression levels and social and **(d)** so preference scores in MOR-HNK mice.

### 3.6 Residual transcriptomic signatures in MOR-HNK compared to saline-treated mice

To assess the extent to which (*2R,6R*)-HNK treatment normalized gene expression profiles in opioid-abstinent mice (MOR-HNK) relative to opioid-naive controls (SAL-SAL), we performed differential expression analysis comparing MOR-HNK *vs.* SAL-SAL groups. Despite evidence of broad transcriptomic recovery, this analysis identified 186 DEGs, including 48 upregulated and 138 downregulated, that remained significantly dysregulated following in MOR-HNK mice compared to opioid-naive controls. Gene ontology analysis of the 48 upregulated DEGs revealed enrichment of immune regulatory processes, including the negative regulation of adaptive immune response, T-helper 17 (Th17) type immune response, and interleukin-10 (IL-10) production (**Figure 7a**). These pathways suggest that immune modulation, particularly involving regulatory and anti-inflammatory signaling, persists in the hippocampus following (*2R,6R*)-HNK treatment, potentially reflecting an ongoing resolution phase of neuroimmune adaptation.

**Figure 7.**
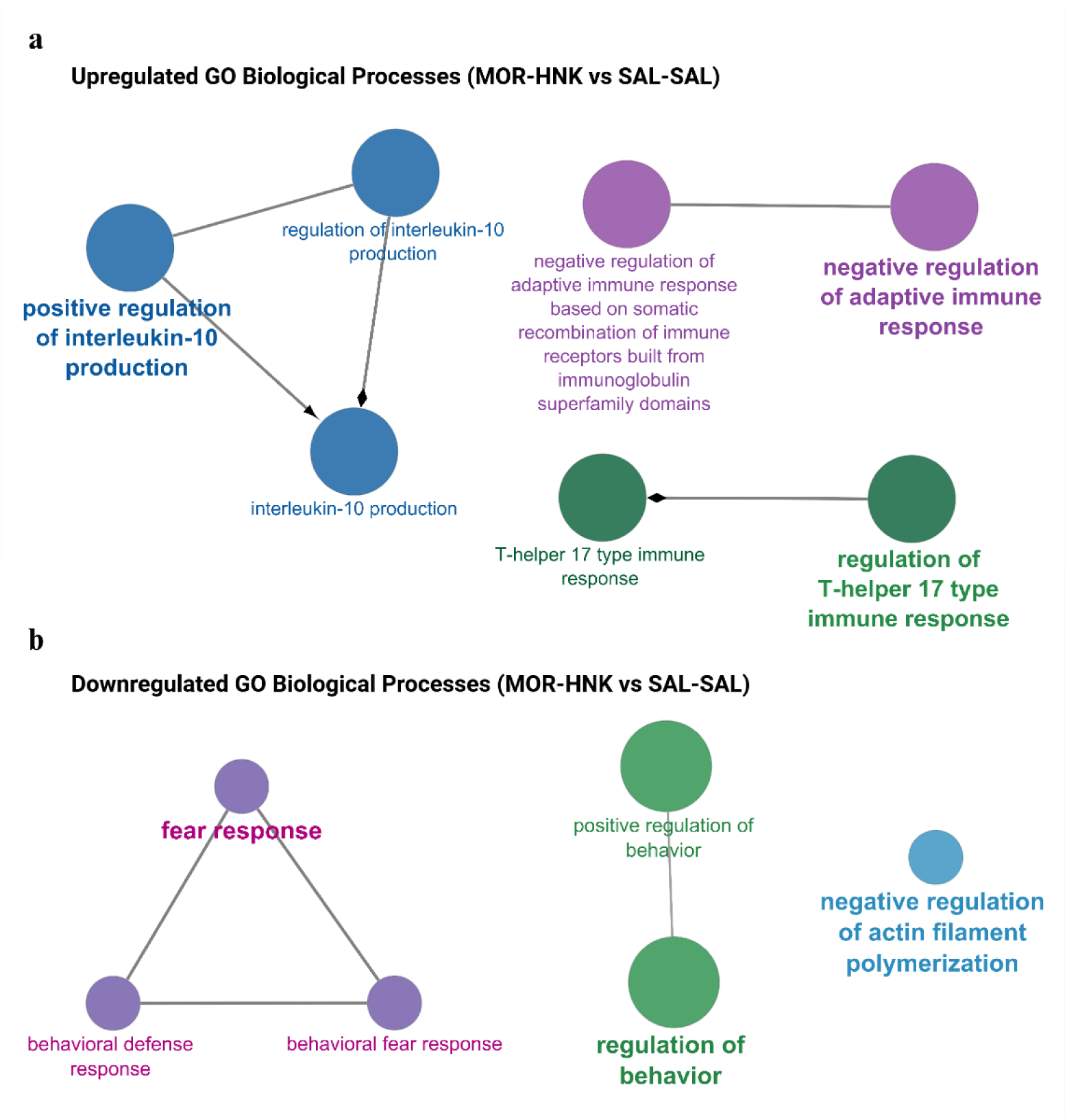
Residual transcriptomic profiles of MOR-HNK vs. SAL-SAL mice. **(a)** GO enrichment analysis of upregulated DEGs in the MOR-HNK vs. SAL-SAL comparison, showing biological processes associated with immune regulation. **(b)** GO enrichment analysis of downregulated DEGs in the MOR-HNK vs. SAL-SAL comparison, illustrating terms related to behavioral regulation and cytoskeletal organization.

Interestingly, biological processes such as the negative regulation of adaptive immune response and T-helper 17 type immune response were also found to be upregulated in the MOR-SAL *vs.* SAL-SAL comparison (see Section 3.2). The persistence of these pathways in the MOR-HNK *vs*. SAL-SAL comparison suggests that (*2R,6R*)-HNK treatment does not fully reverse immune-related transcriptomic alterations induced by chronic morphine exposure. Thus, (*2R,6R*)-HNK may lack the capacity to fully restore immune homeostasis to a naive molecular baseline.

Conversely, analysis of the 138 downregulated DEGs highlighted suppression of GO biological processes related to behavioral regulation and structural dynamics (**Figure 7b**). Enriched terms included regulation of behavior, behavioral defense response, fear response, and positive regulation of behavior, as well as negative regulation of actin filament polymerization. These findings point to residual attenuation of molecular programs associated with behavioral responsiveness and cytoskeletal remodeling, suggesting that (*2R,6R*)-HNK treatment, while restorative, may not fully reinstate all neurobehavioral regulatory pathways to baseline levels.

Together, these results indicate that while (*2R,6R*)-HNK induces substantial recovery of gene expression disrupted by chronic opioid exposure, a distinct subset of immune- and behavior-related pathways remains differentially expressed compared to opioid-naive controls, reflecting an intermediate molecular phenotype.

### 3.7 Standard behavioral assays fail to capture transcriptomic signatures in MOR-HNK mice

To determine whether residual transcriptomic alterations in MOR-HNK mice translated into measurable behavioral differences, we compared sucrose and social preference scores between MOR-HNK vs. SAL-SAL groups. No significant differences were observed in either sucrose (adjusted *p* = 0.7172) or social preference (adjusted *p* = 0.8041) (**Figure 2a, b**), indicating that overt behavioral performance was comparable between groups in these standard assays.

However, the persistent enrichment of immune- and behavior-associated gene pathways in MOR-HNK mice, as indicate by the enrichment results of the upregulated and downregulated DEGs found between MOR-HNK *vs.* SAL-SAL (**Figure 7a, b**), suggests that subtle neurobehavioral alterations may remain undetected by these tests. The data imply that (*2R,6R*)-HNK treatment induces a molecular phenotype that, while broadly normalized, is not identical to the naive state and may underlie latent functional changes beyond the resolution of sucrose and social preference measures. These findings highlight a potential mismatch between transcriptomic sensitivity and behavioral assay specificity, emphasizing the limitations of conventional behavioral models in fully capturing the scope of treatment-induced recovery.

### 3.8 Random Forest ML identifies predictive genes of (*2R,6R*)-HNK treatment response in morphine-exposed mice

To identify genes most predictive of treatment response, we applied a Random Forest classification model using the 200 DEGs identified between MOR-HNK *vs.* MOR-SAL mice. The model achieved an OOB error estimate of 25%, indicating reasonable classification performance despite the relatively small sample size. This suggests that gene expression patterns partially, but not completely, distinguish MOR-HNK-treated animals from morphine-exposed controls at the transcriptomic level.

Variable importance analysis revealed a subset of genes that contributed most strongly to classification accuracy. Based on the MeanDecreaseGini metric, the top 10 most predictive genes were *Cenpk*, *T2*, *Gm17276*, *Il1rapl1*, *Avpr1a*, *Rpph1*, *Cfap52*, *Krtap17.1*, *Ctla2b*, and *Ly9* (**Figure 8**). These genes exhibited the greatest contribution to reducing classification uncertainty and are therefore strong candidate molecular markers of (*2R,6R*)-HNK treatment response.

**Figure 8:**
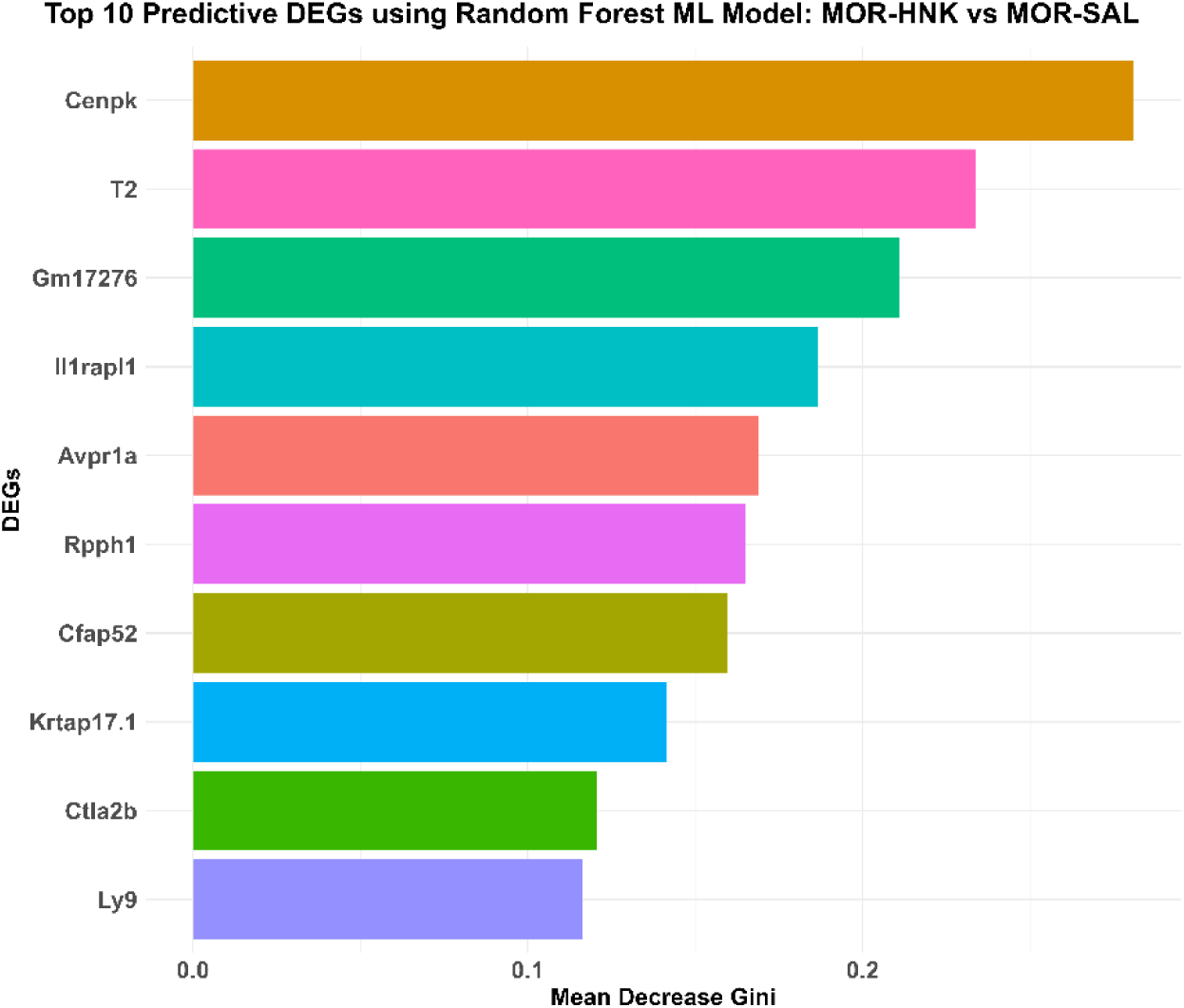
Top 10 predictive DEGs identified by Random Forest classification in the MOR-HNK vs. MOR-SAL comparison. Bar plot shows the top 10 genes ranked by Mean Decrease Gini, reflecting their importance in distinguishing morphine-exposed mice treated with (2R,6R)-HNK (MOR-HNK) from untreated morphine-exposed controls (MOR-SAL).

Importantly, two of the top 10 genes, specifically Interleukin 1 receptor accessory protein-like 1 (*Il1rapl1)* and Cytotoxic T lymphocyte-associated protein 2 beta (*Ctla2b)*, were also among the 55 DEGs identified as exhibiting reversed expression patterns between MOR-SAL *vs*. SAL-SAL and MOR-HNK *vs*. MOR-SAL comparisons. This overlap suggests that these genes may not only mark individual variability but also reflect consistent transcriptional reversal associated with therapeutic recovery. *Il1rapl1*, for instance, is involved in synaptic organization and cognitive function, while *Ctla2b* is linked to immune modulation, reinforcing the relevance of these genes to behavioral and molecular restoration.

Overall, the Random Forest model highlights a focused set of DEGs that best differentiate MOR-HNK-treated animals from morphine-exposed controls, including a subset that also shows population-level reversal, offering particularly promising biomarkers for future mechanistic investigation and validation.

## 4. Discussion

The antidepressant metabolite (*2R,6R*)-HNK reversed morphine-induced upregulation of *Ttr* in a chronic escalating-dose morphine mouse model. Given that protracted opioid abstinence is often accompanied by depressive symptoms, this is a notable finding. *Ttr* is consistently upregulated in the hippocampus in mouse models of stress-induced depression, such as the forced swim stress and chronic social defeat stress (CSDS) paradigms ^51^. Furthermore, viral-mediated *Ttr* overexpression in the hippocampus induces depression-like behaviors and increases proinflammatory gene expression, linking *Ttr* to neuroinflammation, a key mechanism in depression ^51^. Notably, *Ttr* levels were higher in stress-susceptible *vs.* resilient CSDS mice, suggesting specificity to depressive phenotypes ^51^. Supporting this, *Ttr* knockout mice exhibit increased exploratory behavior and reduced depression-like symptoms ^52^. In a recent study utilizing a chronic stress mouse model and *in vivo* epigenome editing, vitamin B12 was shown to promote stress resilience by reducing DNA methylation at the *Ttr* promoter, thereby regulating its expression ^53^. These findings identify *Ttr* as a key epigenetically regulated mediator of stress and depression-related behaviors, with its expression in the prefrontal cortex directly influencing mood outcomes, highlighting *Ttr* as a potential therapeutic target for stress-related disorders ^53^.

Although *Ttr* expression was the most significantly DEGs found to be downregulated and reversed by (*2R,6R*)-HNK treatment, its expression levels did not significantly correlate with either sucrose or social preference, suggesting that *Ttr* regulation may reflect broader physiological adaptations rather than directly mediating behavioral recovery. In the context of addiction neurobiology, *Ttr* is not known to drive ventral hippocampal functions such as reward processing, emotional regulation, or motivational learning, the core behavioral domains disrupted by chronic opioid exposure. Given its primary role as a transporter of thyroxine and retinol in the cerebrospinal fluid, *Ttr* expression changes in the ventral hippocampus likely reflect systemic shifts in brain metabolism, neuroinflammatory tone, or general stress responses rather than direct modulation of addiction-relevant neural circuits.

Similarly, although *Cd5* was the most strongly upregulated DEG among the reversal set, it did not significantly correlate with either social preference or sucrose preference. This lack of association suggests that, despite its robust transcriptional reversal following (*2R,6R*)-HNK treatment, *Cd5* may not contribute directly to behavioral recovery outcomes. Instead, its regulation may reflect broader immune modulation unrelated to the specific behavioral domains assessed in this study. *Cd5* is a transmembrane glycoprotein predominantly expressed on T cells and a subset of B cells known as B-1a cells ^54^. Importantly, *CD5* functions as a negative regulator of T cell receptor (TCR) signaling, helping to dampen immune activation^55^. Its upregulation in MOR-HNK mice may therefore reflect an enhanced immunoregulatory state induced by (*2R,6R*)-HNK, potentially contributing to the broader anti-inflammatory environment observed in the hippocampus. However, given the lack of correlation with behavioral outcomes, *Cd5* may act as a general marker of immune adaptation rather than a gene specifically linked to functional recovery.

These findings highlight the critical importance of integrating transcriptomic profiling with individual-level behavioral analyses to identify molecular changes that are mechanistically relevant to therapeutic efficacy. Indeed, lack of behavioral correlation for a given transcriptomic change does not necessarily imply biological irrelevance. Genes that are reversed or regulated by (*2R,6R*)-HNK but do not associate directly with behavioral outcomes may nonetheless contribute to broader physiological processes, such as modulation of the immune system (e.g *Cd5*), glial activation, or neurovascular regulation. Importantly, although (2R,6R)-HNK altered gene expression in the ventral hippocampus of opioid-naive mice (SAL-HNK), it did not produce significant changes in behavioral assays, including sucrose preference and social preference, compared to saline-treated controls (SAL-SAL). In contrast, robust behavioral improvements were evident when (*2R,6R*)-HNK was administered to mice previously exposed to morphine (MOR-SAL *vs.* SAL-SAL), a model of opioid-induced deficits in reward and social behavior. This pattern indicates that the therapeutic effects of (*2R,6R*)-HNK emerge primarily in a pathological context, specifically, when neural circuits are disrupted by prior drug exposure, while its impact remains minimal under baseline, opioid-naive conditions. This context-dependent efficacy underscores the compound’s potential as a targeted intervention for conditions involving stress, addiction, or neuroplasticity impairments, with a low risk of affecting normal behavioral states.

Transcriptomic profiling revealed persistent alterations in immune- and behavior-associated pathways, indicating that (*2R,6R*)-HNK treatment in opioid-abstinent mice does not fully restore hippocampal gene expression to a naive baseline. The absence of significant behavioral differences between MOR-HNK *vs*. SAL-SAL mice in sucrose and social preference tests may reflect the limited sensitivity of these assays to detect more nuanced or domain-specific behavioral effects. The downregulation of gene sets involved in behavioral regulation, fear response, and actin cytoskeleton dynamics, alongside upregulation of immune modulatory processes such as interleukin-10 signaling and Th17 activity, suggests an intermediate or compensatory molecular state. These patterns may reflect lingering adaptations to chronic opioid exposure or ongoing neurobiological remodeling induced by (*2R,6R*)-HNK. This disconnects between normalized overt behavior and persistent molecular divergence highlights the limitations of standard behavioral assays and the need for more targeted models capable of detecting latent neurobiological states.

The behavioral assays used in this study, sucrose and social preference, are well-established for assessing reward and social behavior in models of opioid exposure and antidepressant treatment ^40,41,43,47,56^. However, the residual transcriptomic alterations observed suggest dysfunction in domains such as fear processing, neuroimmune interaction, and cognitive regulation, which are not fully captured by these measures. Supporting this, studies in post-traumatic stress Disorder (PTSD) models have shown that (*2R,6R*)-HNK reduces fear- and anxiety-like behaviors ^57^. More recently, (*2R,6R*)-HNK was found to specifically alleviate fear-related responses and restore hippocampal Glutamate Ionotropic Receptor AMPA Type Subunit 1 *(GluA1)*, Brain-Derived Neurotrophic Factor *(BDNF)*, and *Nerve Growth Factor Inducible (VGF)* expression when administered during the reconsolidation phase of fear memory ^58^, highlighting a potential mechanism by which this metabolite modulates maladaptive emotional memory. This indicates that (*2R,6R*)-HNK, in the context of morphine abstinence, could downregulate fear response pathways, suggesting its potential suitability for treating comorbid opioid addiction and PTSD.

Given the enrichment of pathways related to fear response, behavioral regulation, and neuroplasticity, future studies should incorporate assays such as fear conditioning, elevated plus maze, and cognitive flexibility tasks (e.g., Y-maze, attentional set-shifting) to better align behavioral assessment with transcriptomic findings. To further evaluate motivational and immune-linked domains, paradigms such as stress-sensitive social preference, progressive ratio reward tasks, and neuroinflammatory models including LPS challenge or wheel-running may help detect subtle functional effects like fatigue, malaise, or latent affective disturbances. Together, these approaches may provide a more comprehensive evaluation of therapeutic recovery following opioid exposure and better capture the functional impact of (*2R,6R*)-HNK at both molecular and behavioral levels.

Furthermore, biological processes such as the negative regulation of adaptive immune response and T-helper 17 type immune response were found to be upregulated in all morphine-abstinence mice, irrespective of their saline or (*2R,6R*)-HNK treatment, compared with saline controls. Chronic opioid exposure is known to dysregulate immune function, often leading to a pro-inflammatory environment within the central nervous system (CNS) ^59^. In particular, chronic morphine administration has been shown to increase the number of circulating regulatory T cells and enhance the functional activity of Th17 cells, contributing to sustained neuroinflammation and altered T cell dynamics ^60^. Th17 cells are a subset of CD4⁺ T cells, that primarily secrete interleukin-17 (Il-17) and play a key role defense against extracellular bacteria and fungi at mucosal surfaces (like lungs, gut, skin) ^61^. Controlled Th17 responses are good for immune defense, but excessive or dysregulated Th17 responses cause inflammation and tissue damage, leading to inflammatory diseases ^62^. Thus, the persistence of Th17-related pathways in morphine-abstinent mice treated with (*2R,6R*)-HNK suggests that treatment does not fully resolve the neuroimmune imbalances initiated by chronic morphine administration or abstinence.

Among the top genes identified by Random Forest as predictive of (*2R,6R*)-HNK treatment response (MOR-HNK *vs.* MOR-SAL), *Il1rapl1* and *Ctla2b* emerged as particularly notable due to their dual relevance: both were among the top 10 most predictive features and were also part of the 55 DEGs identified through comparison of DEGs from the MOR-SAL *vs.* SAL-SAL and MOR-HNK *vs.* MOR-SAL groups. *Il1rapl1* encodes interleukin-1 receptor accessory protein-like 1, which is known to regulate the formation and stabilization of glutamatergic synapses in cortical neurons via the RhoA signaling pathway ^63^. Mutations in *Il1rapl1* have been linked to intellectual disability and autism spectrum disorders ^63^, underscoring its critical role in neurodevelopment and synaptic plasticity. Its reversal and predictive significance suggest that synaptic remodeling processes may be central to the effects of (*2R,6R*)-HNK treatment. In contrast, *Ctla2b* encodes a cytotoxic T lymphocyte-associated protein involved in immune regulation, particularly in T-cell suppression. Its predictive importance supports the idea that normalization of certain neuroimmune pathways, such as those involving CTL regulation, contributes to therapeutic response. Notably, while Ctla2b-related CTL activity appears reversed by HNK, Th17-related pathways, a subset of CD4⁺ T-cell– mediated immunity, remain upregulated in MOR-HNK mice, indicating that (2R,6R)-HNK may only partially restore immune homeostasis. Together, these findings highlight *Il1rapl1* and *Ctla2b* as promising molecular links between transcriptomic plasticity and behavioral restoration in the context of opioid abstinence-induced neuroadaptation.

## 5. Conclusion

This study demonstrates that the antidepressant metabolite (*2R,6R*)-HNK effectively reverses morphine-induced behavioral deficits and associated transcriptomic alterations in the ventral hippocampus, supporting its potential as a context-specific therapeutic agent. Key molecular targets such as *Ttr* and *Cd5* showed robust regulation by (*2R,6R*)-HNK, though their expression did not directly correlate with behavioral recovery, highlighting the complexity of disentangling mechanistic vs compensatory molecular changes. Notably, (*2R,6R*)-HNK induced transcriptional changes even in opioid-naive mice without altering behavior, reinforcing its selective efficacy in pathological states such as opioid-induced dysfunction. Persistent immune and neuroplasticity-related transcriptomic changes suggest ongoing remodeling in morphine-abstinent mice, which standard behavioral assays may not fully capture. These findings underscore the need for more nuanced behavioral models and integrated transcriptomic-behavioral analyses to assess therapeutic outcomes. Overall, (*2R,6R*)-HNK exhibits promise as a targeted intervention for opioid-related and stress-associated disorders, with minimal impact on normal brain function.

## Author Contributions

**A. Onisiforou**: Conceptualization, data curation, formal analysis, methodology, writing—original draft, review, and editing. **M. Koumas:** Investigation, writing—review and editing. **A. Michael:** Investigation, writing—review and editing. **P. Zanos:** Conceptualization, investigation, funding acquisition, project administration, writing—review and editing.

## Acknowledgments

The authors would like to thank Dr. Christiana Neophytou and Prof. Panagiotis Papageorgis of the European University Cyprus for providing laboratory space for RNA extraction. We also gratefully acknowledge Dr. Patrick Morris and Dr. Craig Thomas from the National Center for Advancing Translational Sciences (NCATS), NIH, USA, for characterizing and supplying (2R,6R)-HNK.

## Data Availability Statement

The data supporting the findings of this study are available from the corresponding authors upon reasonable request.

